# A Minimal Plasma Proteome-Based Biomarker Panel for Accurate Prostate Cancer Diagnosis

**DOI:** 10.1101/2025.11.05.686712

**Authors:** Syed Ahsan Shahid, Ahmed Al-Harrasi, Adil Al-Siyabi

**Author notes:** Corresponding author: **Adil Al-Siyabi**, Assistant Professor, Natural and Medical Sciences Research Center, University of Nizwa, Birkat Al-Mouz, Nizwa, 616, Oman.

## Abstract

Early and accurate diagnosis of prostate cancer (PRC) remains a major clinical challenge, particularly with existing biomarker panels relying on invasive sampling or large biomarker panels with limited interpretability. Here, we present a machine learning framework for discovering compact and biologically grounded plasma protein signatures for PRC classification using publicly available pan-cancer proteomic data. We coupled a genetic algorithm-based protein identification method with LASSO-regularized logistic regression to identify minimal protein subsets optimized for diagnostic performance. A 14-protein panel, recurrent across 1,000 genetic algorithm iterations, achieved a mean accuracy of 98.0%, an F1 score of 0.98, and an ROC AUC of 0.997 on a held-out test dataset. This performance exceeded models trained on high dimensionality data (>1,400 proteins) and surpassed published transcriptomic, methylomic, and cfDNA classifiers, many of which reported AUCs less than 0.91. Functional analysis revealed enrichment in protease binding and DNA repair pathways, with known markers such as beta-microseminoprotein (MSMB) and poly(ADP-ribose) polymerase 1 (PARP1) appearing alongside under-characterized proteins like IGSF3 and XG. Models trained only on previously reported PRC-associated proteins showed lower performance, highlighting the added diagnostic value of including novel, data-driven candidates. This study outlines a scalable proteomic workflow and demonstrates that high diagnostic performance can be achieved using small, interpretable panels derived from blood-based proteomics. The findings lay the foundation for the development of interpretable, clinically deployable assays for PRC detection and risk stratification.

**Graphical abstract:** 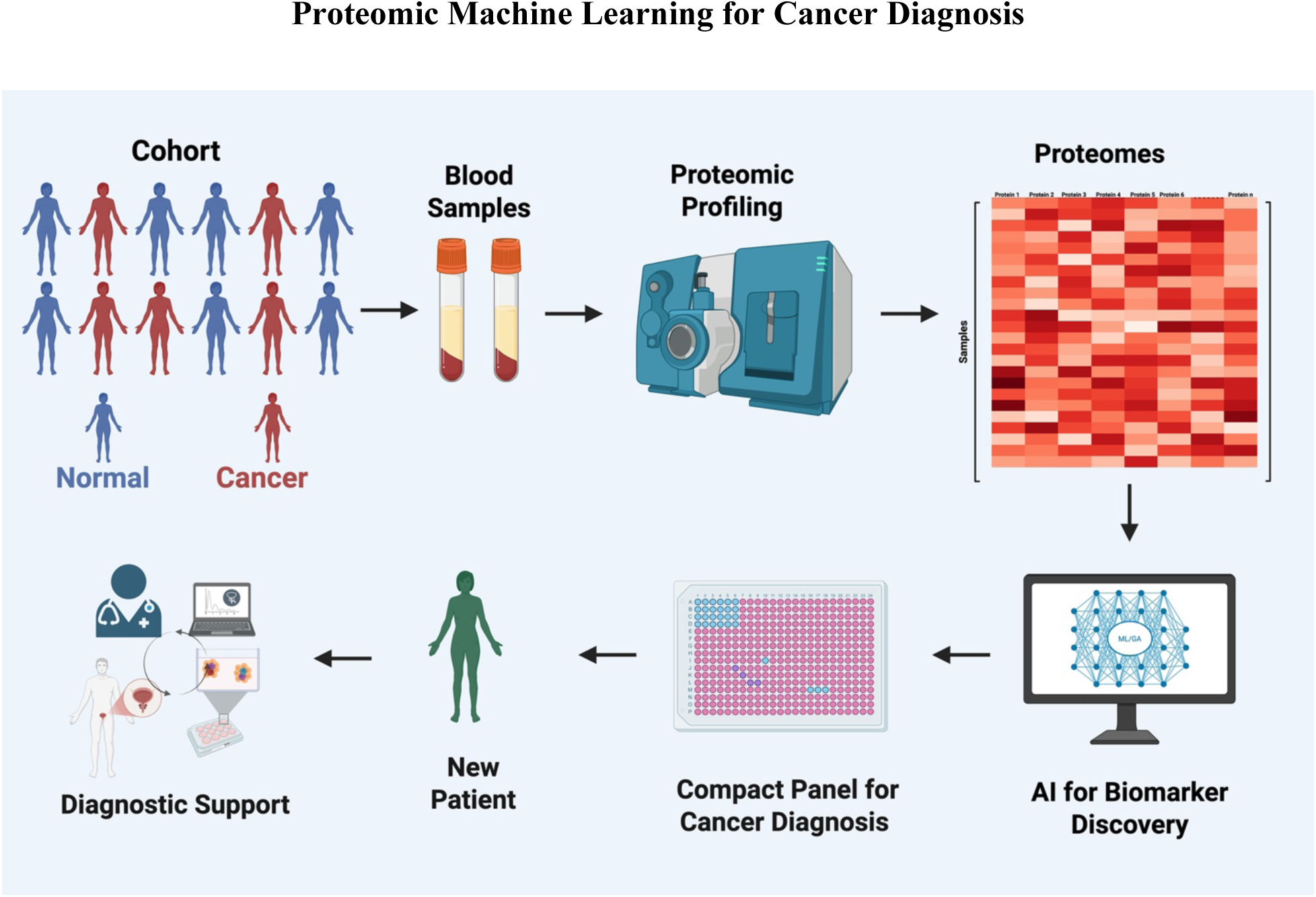

Overview of the study workflow: blood-derived proteomic profiles from cancer and normal cohorts were analyzed using machine learning to identify a compact diagnostic panel, enabling clinical decision support for new patients.

## Introduction

Prostate cancer is one of the most commonly diagnosed malignancies among men worldwide and remains a major contributor to cancer-related mortality [1]. According to the Global Cancer Observatory (GLOBOCAN) and Global Burden of Diseases (GBD) reports, prostate cancer accounted for over 1.4 million new cases and approximately 375,000 deaths globally in 2020 [2, 3]. It is the leading cancer among men in more than 100 countries and ranks among the top two cancers in terms of incidence across nearly every continent [1]. The burden of prostate cancer continues to rise, largely driven by aging populations, lifestyle changes, and improved diagnostic access in high-income and transitional economies [4]. Beyond its epidemiological footprint, prostate cancer significantly affects the physical, emotional, and social well-being of patients, particularly when diagnosed at advanced stages, where therapeutic options are limited and the risk of metastasis and mortality is higher [5]. It has been reported by the Lancet Commission on Prostate Cancer that there will be a significant annual increase in new prostate cancer cases from 1.4 million in 2020 to 2.9 million by 2040 [6].

Early detection plays a pivotal role in reducing the burden of prostate cancer. When diagnosed at localized stages, prostate cancer is highly treatable, with five-year survival rates exceeding 95% [7]. However, survival drops dramatically once the disease metastasizes, highlighting the importance of timely and accurate diagnosis. Early detection not only improves patient prognosis but also helps avoid overtreatment, allowing clinicians to distinguish between indolent and aggressive disease [6]. Current screening practices, which primarily involve prostate-specific antigen (PSA) testing and digital rectal examination, have improved early detection rates but are fraught with limitations. PSA testing lacks the specificity required to distinguish malignant from benign conditions such as prostatitis or benign prostatic hyperplasia (BPH), leading to high false-positive rates and unnecessary biopsies [8]. Moreover, PSA levels do not provide insight into the molecular characteristics or aggressiveness of tumors, making it difficult to tailor treatment decisions based on underlying biology [9].

To address these limitations, there has been growing interest in the use of blood-based proteomic profiling as a non-invasive diagnostic tool [10]. Blood is an attractive medium for biomarker discovery because it can be obtained with minimal discomfort and reflects both systemic and tumor-specific alterations [11]. Advances in high-throughput proteomic technologies, particularly the proximity extension assay (PEA), have enabled the simultaneous quantification of over a thousand proteins in small volumes of plasma with high sensitivity and specificity [12]. This has opened new avenues for identifying cancer-specific protein signatures and monitoring disease progression, treatment response, or recurrence. The integration of proteomics into the diagnostic landscape has the potential to transform clinical workflows by enabling liquid biopsy-based assays that are both minimally invasive and rich in molecular information [13].

Despite these technological advancements, interpreting proteomic data for clinical application remains challenging. Proteomic datasets are inherently high-dimensional, with thousands of variables (proteins) measured across a limited number of samples [14]. This “large p, small n” scenario complicates traditional statistical analysis and increases the risk of overfitting, especially when building predictive models [15, 16]. Furthermore, the complexity of biological systems means that disease signatures often involve intricate interactions among proteins, rather than isolated markers [17]. As such, sophisticated computational methods are required to uncover meaningful patterns and reduce dimensionality without losing predictive power.

Machine learning (ML) has emerged as a powerful tool for analyzing complex biomedical data. ML algorithms are capable of handling nonlinear relationships, interactions, and noise in high-dimensional datasets, making them well-suited for proteomic biomarker discovery [18]. In cancer research, ML has been applied to identify diagnostic, prognostic, and predictive biomarkers from a variety of omics data types, including genomics, transcriptomics, and proteomics [19]. ML approaches such as logistic regression (LR), support vector machines (SVM), random forests (RF), and regularized regression models have shown promise in classifying cancer subtypes and predicting treatment outcomes [20]. However, these models often require extensive feature engineering, hyperparameter tuning, and rely on large feature sets, which limits their interpretability and hinders clinical implementation. Moreover, the selection of informative biomarkers is often carried out manually or using suboptimal methods, reducing reproducibility and scalability [21–23].

To overcome these limitations, evolutionary algorithms such as genetic algorithms (GAs) have been proposed for automated and optimized feature selection. GAs mimic the process of natural selection to identify subsets of features that maximize model performance while minimizing redundancy [24]. When combined with ML frameworks, GAs offer a systematic and scalable approach for identifying minimal, high-performance biomarker panels [25]. This is particularly important for clinical applications, where simpler models with fewer features are more likely to be adopted due to reduced cost, complexity, and regulatory burden.

In this study, we present a GA-based ML pipeline for the discovery of blood-derived protein signatures specific to prostate cancer. Using publicly available proteomic data from a large pan-cancer cohort profiled with PEA [12], we focused on prostate cancer samples to develop compact, high-accuracy diagnostic models. Our approach iteratively evolved protein subsets and evaluated their predictive performance across bootstrapped splits, enabling both model robustness and feature selection stability. We further assessed the biological relevance of the identified proteins through enrichment analyses, aiming to connect the selected biomarkers to known or emerging pathways involved in prostate tumorigenesis.

By integrating blood-based proteomics, evolutionary feature selection, and ML classification, our framework seeks to address key challenges in prostate cancer diagnostics, namely, the need for accurate, interpretable, and scalable biomarker panels. The resulting models not only achieve high diagnostic performance, accuracy: 98.0%, ROC AUC: 0.997 with a minimal number of features (14 proteins) but also offer insights into the biological underpinnings of prostate cancer. These findings lay the groundwork for future efforts to develop clinically deployable assays for early detection and personalized risk stratification in prostate cancer management.

## Methodology

### Dataset Source and Preprocessing

Plasma proteomic data were obtained from a publicly available pan-cancer cohort profiled using the Olink Explore 1536 platform [12]. This platform combines proximity extension assay (PEA) with next-generation sequencing (NGS), enabling highly sensitive and multiplexed quantification of over 1,400 plasma proteins from minimal sample volumes. Each sample was accompanied by detailed clinical annotations, including diagnosed cancer type, allowing disease-specific stratification.

For this study, all available prostate cancer (PRC) samples were extracted and compared against a balanced control cohort derived from other tumor types. To minimize sampling bias and prevent overrepresentation of any single cancer type, non-PRC samples were stratified by cancer type, and equal numbers were randomly selected from each group to match the number of PRC cases. This strategy ensured comparable group sizes and balanced class representation across analyses. All downstream preprocessing, feature selection, and modeling were performed using this curated, balanced dataset, providing a robust foundation for subsequent machine learning and statistical evaluations.

### Genetic Algorithm Optimization for Protein Selection

To identify compact and high-performing plasma protein panels for prostate cancer (PRC) classification, a genetic algorithm (GA)-based protein selection strategy was implemented using the *sklearn-genetic* framework. GA is a population-based optimization method that iteratively evolves protein subsets by maximizing predictive performance through simulated natural selection.

The GA was initialized with a population of 1,000 randomly generated protein sets, each constrained to include a maximum of 40 proteins. Selection pressure was applied using ROC AUC as the fitness function, optimized over 300 generations with 4-fold cross-validation. Mutation and crossover probabilities were set empirically at 0.1 and 0.9, respectively. Both logistic regression (L1-penalized) and SVM classifiers were tested, with logistic regression selected as the final model based on interpretability and consistent performance across bootstrap replicates.

The GA process was repeated across 1,000 independent iterations. For each iteration, the dataset was rebalanced to include equal numbers of PRC and non-PRC samples, randomly sampled across tumor types to prevent class or tissue-type bias. The best-performing panel from each run was recorded, along with the fitness trajectory and the selected protein set.

Two complementary outputs were generated, i.e., iteration-specific protein sets, representing the best-performing protein subset per run, and a ranked protein list, quantifying protein selection frequency across all 1,000 iterations.

To evaluate protein set stability, protein occurrence frequencies were used for ranking. These top-ranked proteins were then cumulatively tested using bootstrap-aggregated LASSO logistic regression.

### Machine Learning Model Training and Evaluation

To evaluate the diagnostic performance of candidate protein subsets, we implemented a LASSO-regularized logistic regression framework with bootstrap aggregation and incremental feature inclusion. Protein selection was guided by GA derived sets according to each analytical context. Model training and evaluation were performed using the *LogisticRegressionCV* class from scikit-learn, with L1 regularization *(penalty=’l1’)* and the *liblinear* solver. Each model was trained on a balanced dataset composed of all PRC samples and an equal number of randomly sampled non-PRC cases. Stratified sampling ensured equal representation across non-PRC cancer types.

For each feature set, we performed 100 bootstrap replicates. In each replicate, samples were resampled with replacement, split into 80% training and 20% test sets (stratified by label), and standardized using Z-score normalization.

Models were evaluated using 5-fold cross-validation within the training set. Performance was assessed on the held-out test data in each replicate. Classification metrics included accuracy, ROC AUC, class-specific precision and recall, F1 scores, predictive values, and macro-averaged F1. Confusion matrices were used to compute class-specific metrics, and results were averaged across bootstrap replicates. Variability was captured using standard deviation estimates for each metric. This approach enabled systematic performance profiling of protein subsets of increasing size and provided stable estimates of classification robustness. All results were saved in tabular format and used to identify the optimal feature panel based on diagnostic performance plateau.

### Differential Expression Analysis for up- and down-regulated proteins

We applied the Limma package to identify proteins differentially expressed between PRC and non-PRC groups. P-values were adjusted using the Benjamini-Hochberg method to control for false discovery rate. Proteins with adjusted p < 0.05 and |log₂ fold change| > 0.5 were considered significant. The top 20 ranked proteins based on adjusted p-values were selected for initial model development.

## Results

### Genetic Algorithm-Driven Protein Selection and Systematic Evaluation of the Protein Panel Performance

#### Iteration-specific optimization and model performance

To identify compact protein panels optimized for classification performance, we implemented a genetic algorithm (GA)-based protein selection strategy. The GA was executed across 1,000 independent iterations. In each iteration, a balanced dataset of PRC and non-PRC samples was generated, and the algorithm searched for subsets of 40 proteins that maximized classification performance. Protein selection was performed separately using logistic regression (L1-penalized) and support vector machine classifiers, with logistic regression selected for final downstream analysis based on overall performance and interpretability. The GA produced two key outputs: (1) iteration-specific protein sets representing the highest-scoring subset from each individual run, and (2) a ranked list of proteins based on their recurrence frequency across all 1,000 iterations.

We evaluated the diagnostic performance of the top 100 protein sets identified across runs by training LASSO logistic regression models on each set. For each protein panel, we ran 100 independent bootstrap-aggregated, five-fold cross-validation to evaluate classification accuracy, ROC AUC, F1 scores, and class-specific precision and recall for both PRC and non-PRC classes (Supplementary Data 1). On average, the best-performing panels achieved an accuracy of 94.4%, F1 scores 87.6%, and ROC AUC of 0.980. Approximately 90% of the top-ranked sets achieved ROC AUCs above 0.95 and F1 scores above 0.80. These panels consistently balanced predictive performance between the PRC and non-PRC classes. Standard deviations across the protein sets remained low, indicating stable performance. For example, top sets showed a standard deviation in ROC AUC under 0.01 and F1 standard deviations around 0.03-0.07. Panels with >90% accuracy maintained F1 standard deviations below 0.06 for both classes, supporting reproducibility (Fig. 2A; Supplementary Data 1).

**Figure 1:**
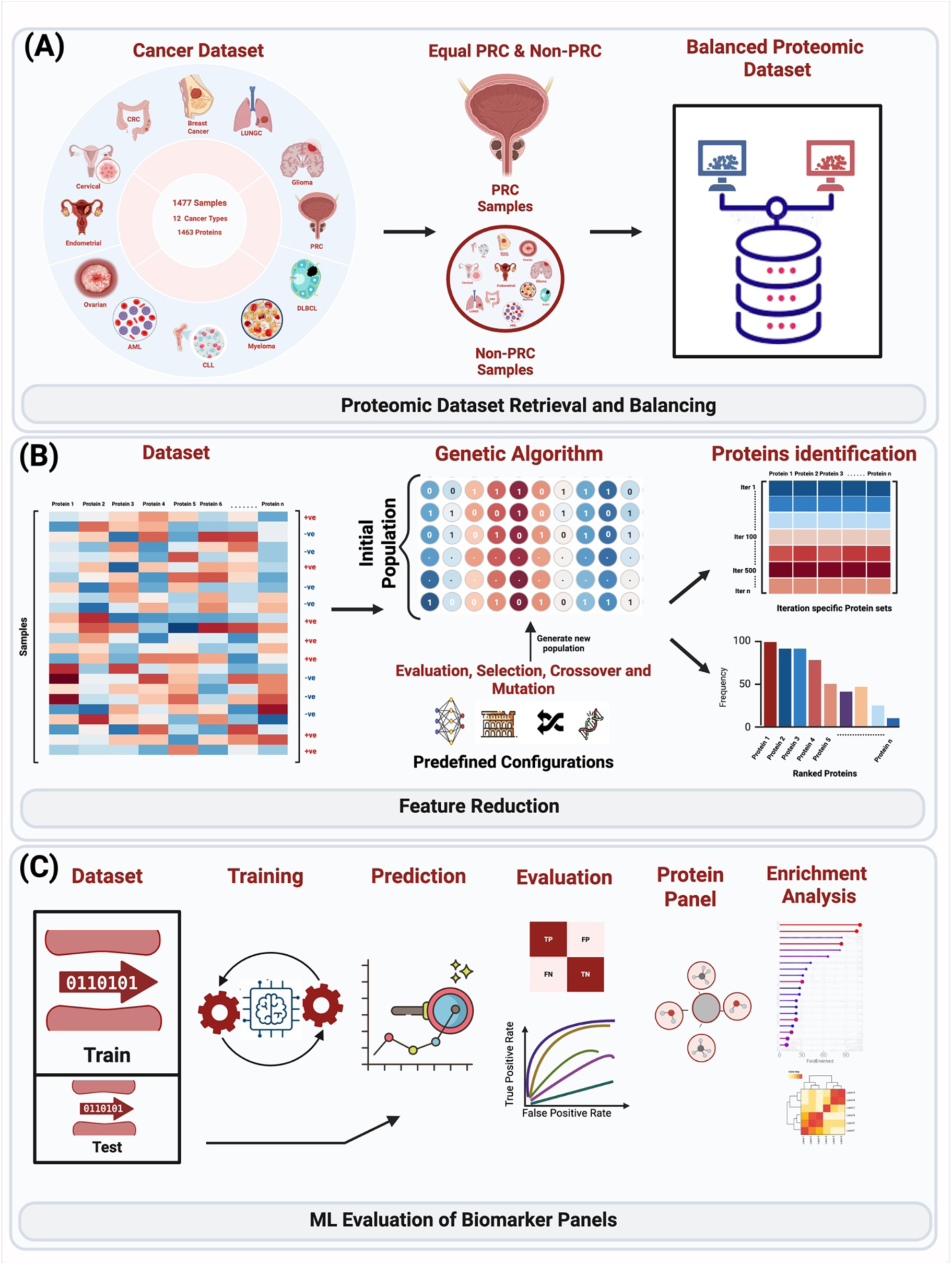
Workflow for proteomic biomarker discovery and evaluation in prostate cancer (PRC). (A) Proteomic dataset retrieval and balancing. A multi-cancer proteomic dataset comprising 1,477 samples across 12 cancer types (1463 proteins) was collected. To avoid bias, equal numbers of PRC and non-PRC samples were selected to generate a balanced proteomic dataset. (B) Feature reduction using a genetic algorithm. The dataset was processed through a genetic algorithm framework, where initial protein subsets underwent evaluation, selection, crossover, and mutation to generate new populations. Iteration-specific protein sets were identified, and proteins were ranked according to selection frequency across runs. (C) Machine learning (ML) evaluation of biomarker panels. The balanced dataset was split into training and test cohorts. ML models were trained and evaluated for predictive performance (ROC, sensitivity, specificity, and confusion matrix). The resulting top-performing protein panels were further assessed by enrichment analysis to determine their biological and clinical relevance.

**Figure 2.**
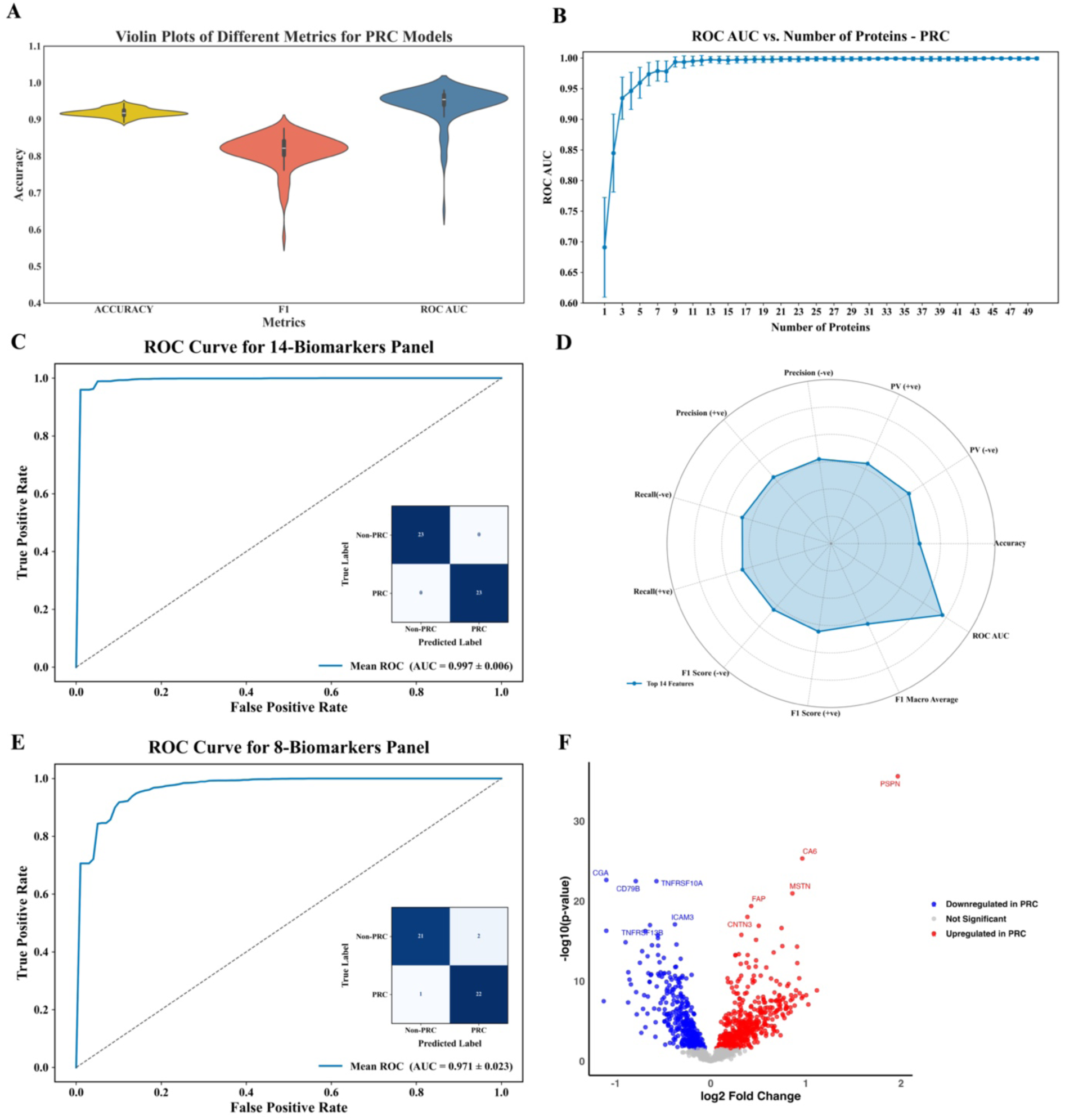
Classification performance evaluation of the identified protein panels. **(A)** Violin plots of accuracy, F1 score, and ROC AUC distributions across 100 bootstrap replicates for the top 100 iteration-specific protein panels. The narrow distributions highlight the stability and reproducibility of the classification performance. **(B)** ROC AUC values of the LASSO logistic regression models plotted against the number of top-ranked proteins used as subsets. Error bars represent standard deviation across 100 bootstrap replicates. A sharp increase in ROC AUC is observed up to 14 proteins, after which performance plateaus. **(C)** This plot presents the ROC curve and its confusion matrix for the 14-protein panel. **(D)** Radar plot showing average classification metrics for the 14-protein panel model across 100 replicates. Metrics include class-specific precision, recall, F1 scores, predictive values, accuracy, macro-averaged F1, and ROC AUC. The plot demonstrates balanced and high performance across both PRC-positive and PRC-negative classes. **(E)** This plot shows the ROC curve for the 8-protein panel and its confusion matrix. **(F)** The Volcano plot shows results from Limma differential expression analysis comparing PRC samples to a balanced set of non-PRC cancer types. Each point represents a plasma protein quantified by PEA. The x-axis indicates log2 fold change, and the y-axis shows -log10 adjusted p-values (Benjamini-Hochberg correction). Proteins significantly upregulated in PRC are shown in red, and downregulated proteins in blue (adjusted p < 0.05). Some of the proteins with high fold changes and strong statistical significance are labeled.

Importantly, some panels achieved near-optimal classification with as few as 23-30 proteins, suggesting that compact sets can provide reliable diagnostics. While several larger panels also performed well, they offered only marginal improvements, reinforcing the efficiency of smaller, GA-optimized subsets. One additional observation was that, although many iteration-specific protein sets yielded classification accuracies above 90%, they often showed lower precision and recall, particularly for one of the classes. This indicates that while the overall accuracy was high, class-specific discrimination was sometimes suboptimal, reinforcing the need for performance-balanced selection criteria beyond accuracy alone. These results confirm that GA effectively identifies concise, high-performing protein signatures for prostate cancer classification.

#### Incremental ranking and identification of the optimal 14-protein panel

In addition to evaluating individual protein sets, we also analyzed the ranked recurrence of proteins across all GA iterations to derive a stable feature importance profile. To evaluate the diagnostic value of the most consistently selected proteins, we ranked all proteins based on how frequently they appeared in the top-performing sets across 1,000 GA iterations. Using this ranked list, we constructed a series of protein subsets starting from the top 1 protein and incrementally adding one protein at a time up to the top 50. Classification performance was evaluated using cumulatively increasing subsets, from the top 1 to the top 50 ranked proteins. For each subset size (1 to 50 proteins), we trained LASSO logistic regression models using 100 bootstrap replicates, each evaluated with five-fold cross-validation. Performance metrics included accuracy, ROC AUC, class-specific precision, recall, and F1 score, all performed on a held-out test set.

Consistent improvements in classification performance were observed as more top-ranked proteins were incorporated. Accuracy rose from 65.8 % with a single top-ranked protein to 98.7 % with 50 proteins. ROC AUC followed a similar trend, increasing from 0.69 to 0.99. F1 scores also improved, reaching 0.98 for the largest panels. Importantly, inter-replicate variability declined with subset size: ROC AUC standard deviation dropped from ∼0.08 with 1 protein to < 0.003 beyond 25 proteins, underscoring model stability with more robust panels (Fig. 2B; Supplementary Data 1).

The panel comprising the top 14 most-repeated proteins demonstrated near-optimal classification with minimal performance gains beyond this point. This subset achieved an average accuracy of 98.0 %, F1 of 0.980, and ROC AUC of 0.997. Class-specific precision and recall were well balanced, with F1 scores of 0.981 (PRC) and 0.979 (non-PRC) (Table 1), and standard deviations below 0.02 across all metrics. Notably, beyond 14 proteins, accuracy and ROC AUC curves plateaued (Fig. 2C), indicating diminishing returns from adding further proteins. Radar plots confirmed uniform distribution of performance across all evaluated metrics for the 14-protein model (Fig. 2D). We also trained classification models using only the top eight most recurrent proteins for comparison. This panel yielded lower classification performance (accuracy 91.3 %, ROC AUC 0.97, F1 0.9) compared to the 14-protein panel (Fig. 2E; Table 1). In contrast to the GA iteration-specific panels, the ranked-protein approach yielded high metric scores, highlighting the 14-protein panel as an efficient and robust diagnostic signature capable of maintaining high performance while minimizing panel redundancy. The distribution of the top 14 recurrent proteins and their structured co-selection patterns are shown in Figure 3A-B, highlighting both individual marker robustness and inter-protein associations.

**Figure 3.**
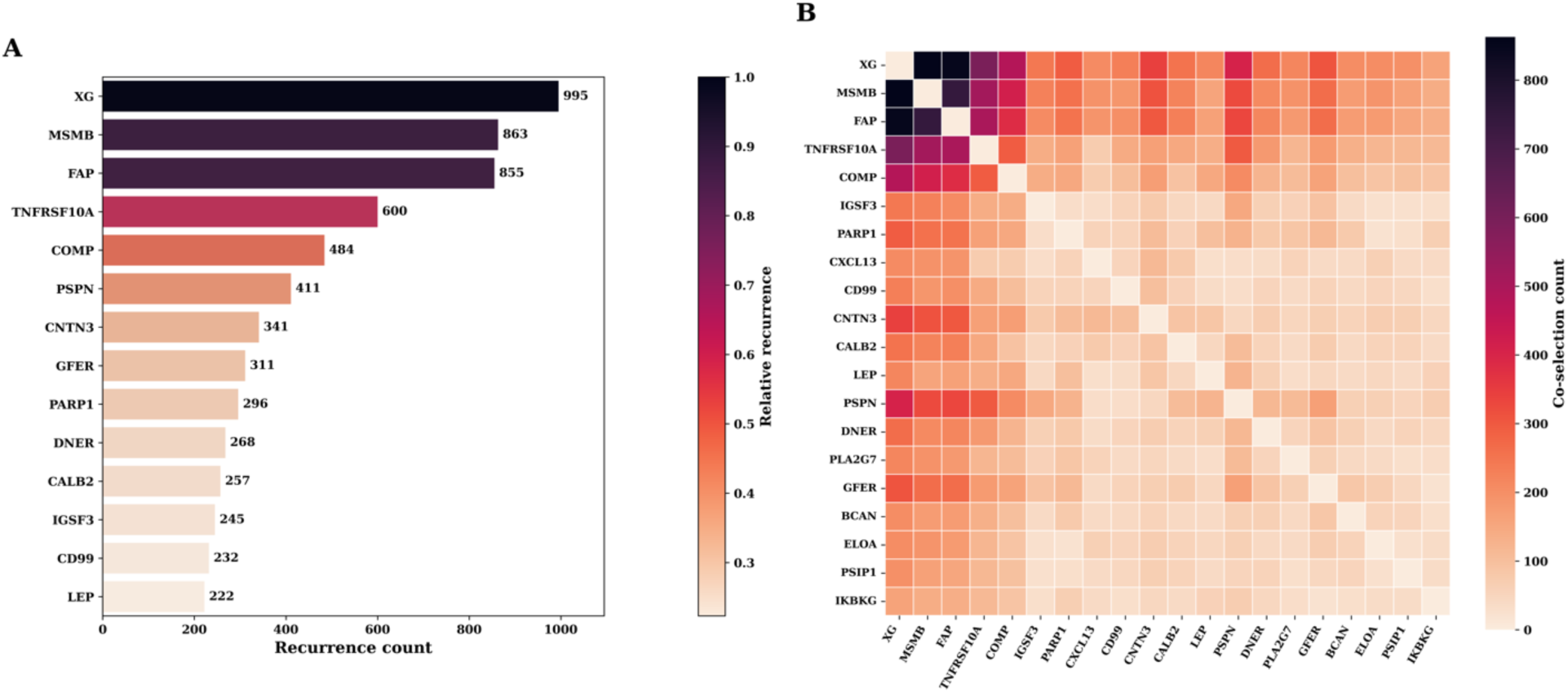
Genetic algorithm-driven protein selection. **(A)** Barplot of the top-most recurrent proteins identified across 1,000 GA iterations. Bar length denotes recurrence count, and color intensity represents relative recurrence frequency. These consistently selected proteins highlight robust features favored by the evolutionary search process. **(B)** Co-selection heatmap of the top 20 recurrent proteins, showing pairwise co-occurrence frequencies across GA-selected feature sets. Warmer colors indicate higher co-selection counts, revealing structured patterns of protein co-selection and potential functional clustering.

**Table 1:** Comparative performance of diagnostic protein panels identified via genetic algorithm (GA), and previously published panels by Alvez et al., 2023 [12] and DEA.

| Panels | Accuracy | Predictive Value (Non-PRC) | Predictive Value (PRC) | Precision (Non-PRC) | Precision (PRC) | Recall (Non-PRC) | Recall (PRC) | F1 score | ROC AUC |
| --- | --- | --- | --- | --- | --- | --- | --- | --- | --- |
| 14 Proteins GA | 0.982 | 0.984 | 0.982 | 0.981 | 0.982 | 0.984 | 0.984 | 0.982 | 0.998 |
| 8 Proteins GA | 0.913 | 0.933 | 0.901 | 0.890 | 0.901 | 0.933 | 0.935 | 0.912 | 0.971 |
| 14 Proteins Alvez et al., 2023 [12] | 0.768 | 0.787 | 0.761 | 0.737 | 0.761 | 0.787 | 0.799 | 0.766 | 0.852 |
| 30 Proteins Alvez et al., 2023 [12] | 0.919 | 0.943 | 0.903 | 0.893 | 0.903 | 0.943 | 0.945 | 0.918 | 0.958 |
| DEA top 20 Proteins | 0.923 | 0.939 | 0.913 | 0.904 | 0.913 | 0.939 | 0.942 | 0.923 | 0.969 |

#### Comparison to Differential Expression-Based Protein Sets

To benchmark our GA-based diagnostic panel, we compared it against protein sets derived from differential expression analysis (DEA) and to previously published panels. In a recently published work [12], the authors implemented a classification approach based on glmnet-regularized regression and achieved above 90% accuracy only when training the model on the full panel of 1,463 proteins. However, performance declined substantially as the number of proteins was reduced. The top 14 proteins identified only achieved an ROC AUC of 0.852.

We applied our LASSO-based classification framework to both their 14- and 30-protein panels. While the 30-protein set showed moderate performance improvements, our GA-optimized 14-protein panel still outperformed both, achieving an accuracy of 98.0%, ROC AUC of 0.997, and F1 of 0.980. Furthermore, we also performed Limma-based DEA on the same dataset (Fig. 2F) and trained a LASSO model on the top 20 differentially expressed proteins. This model achieved high classification performance (accuracy: 92.3%, ROC AUC: 0.969, F1: 92.6%), but still fell short of our GA-derived panel. These comparisons highlight that both our protein selection method (GA) and classification approach (bootstrap-aggregated LASSO) yield superior diagnostic performance compared to traditional DEA and glmnet-based models. They also show that optimal performance does not require large panels, reinforcing the utility of compact, biologically informed protein signatures (Table 1).

#### Biological validation and literature-supported interpretation of selected proteins

To assess the biological plausibility of the final GA-selected 14-protein panel, we conducted a structured literature survey. Each protein was annotated for its known molecular functions, prior evidence in PRC or other cancers, and putative biological roles (Table 2). The resulting panel includes a combination of well-studied PRC-associated proteins and several less-characterized candidates, offering both validation and novel hypotheses.

**Table 2:**
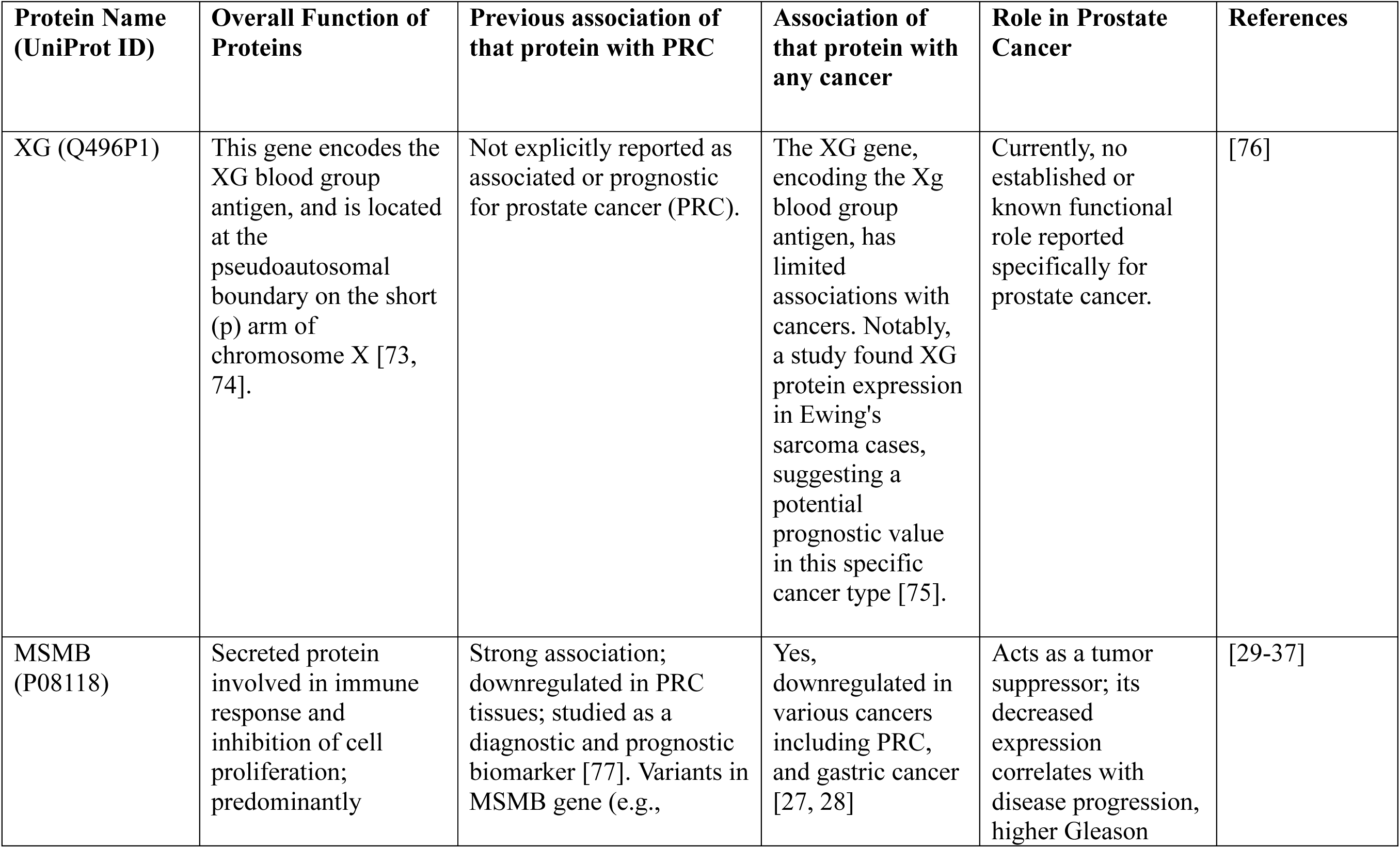
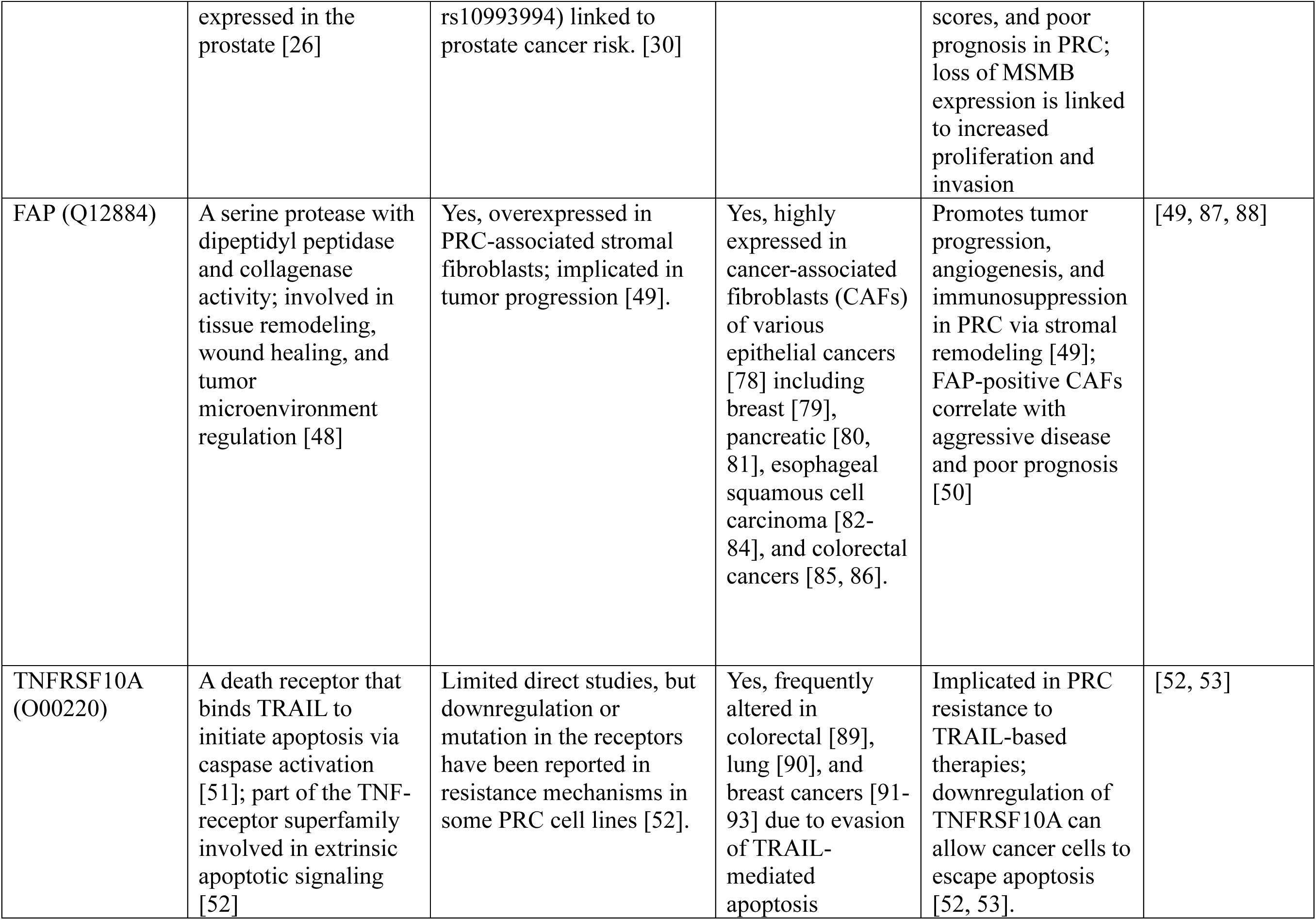
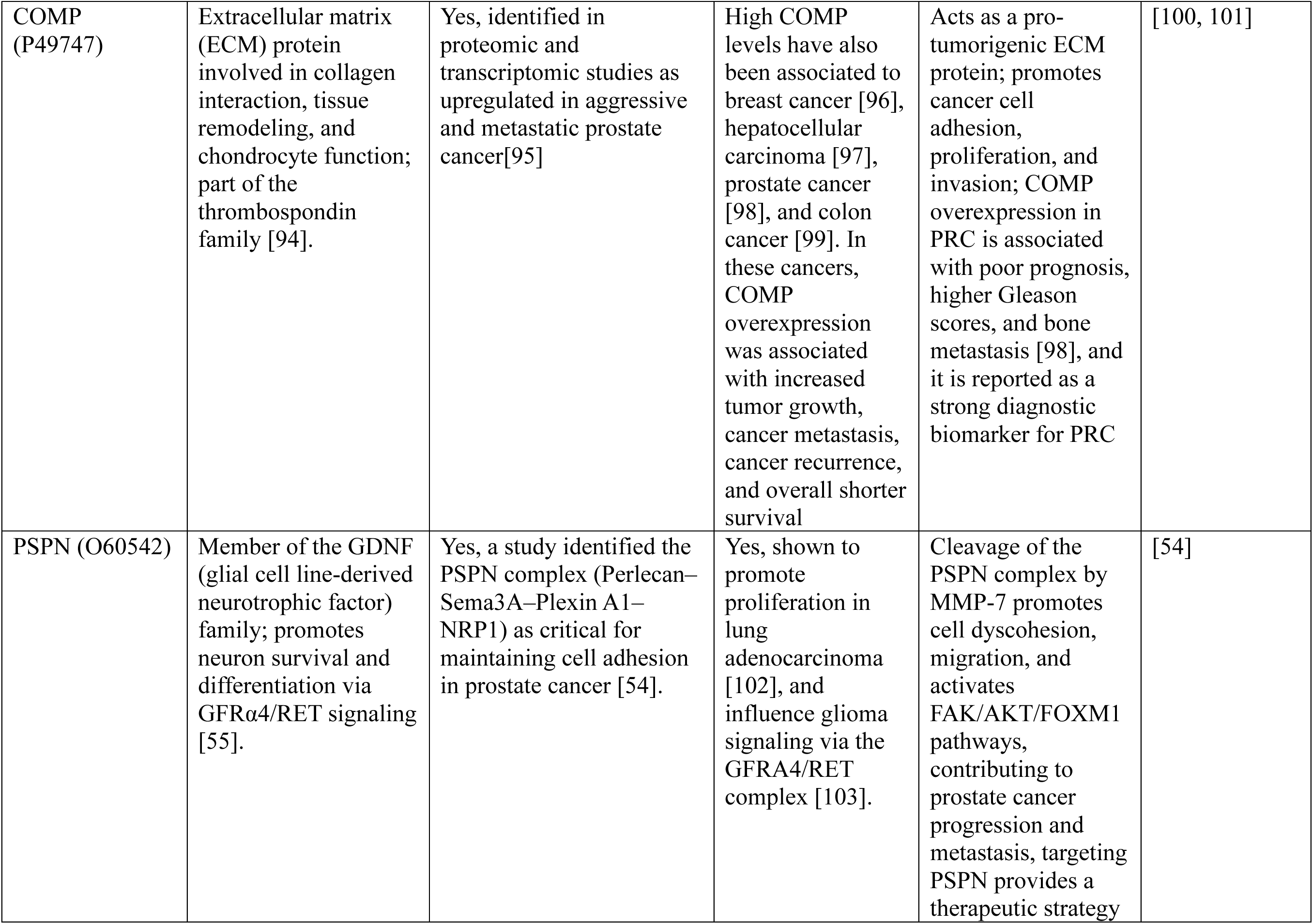
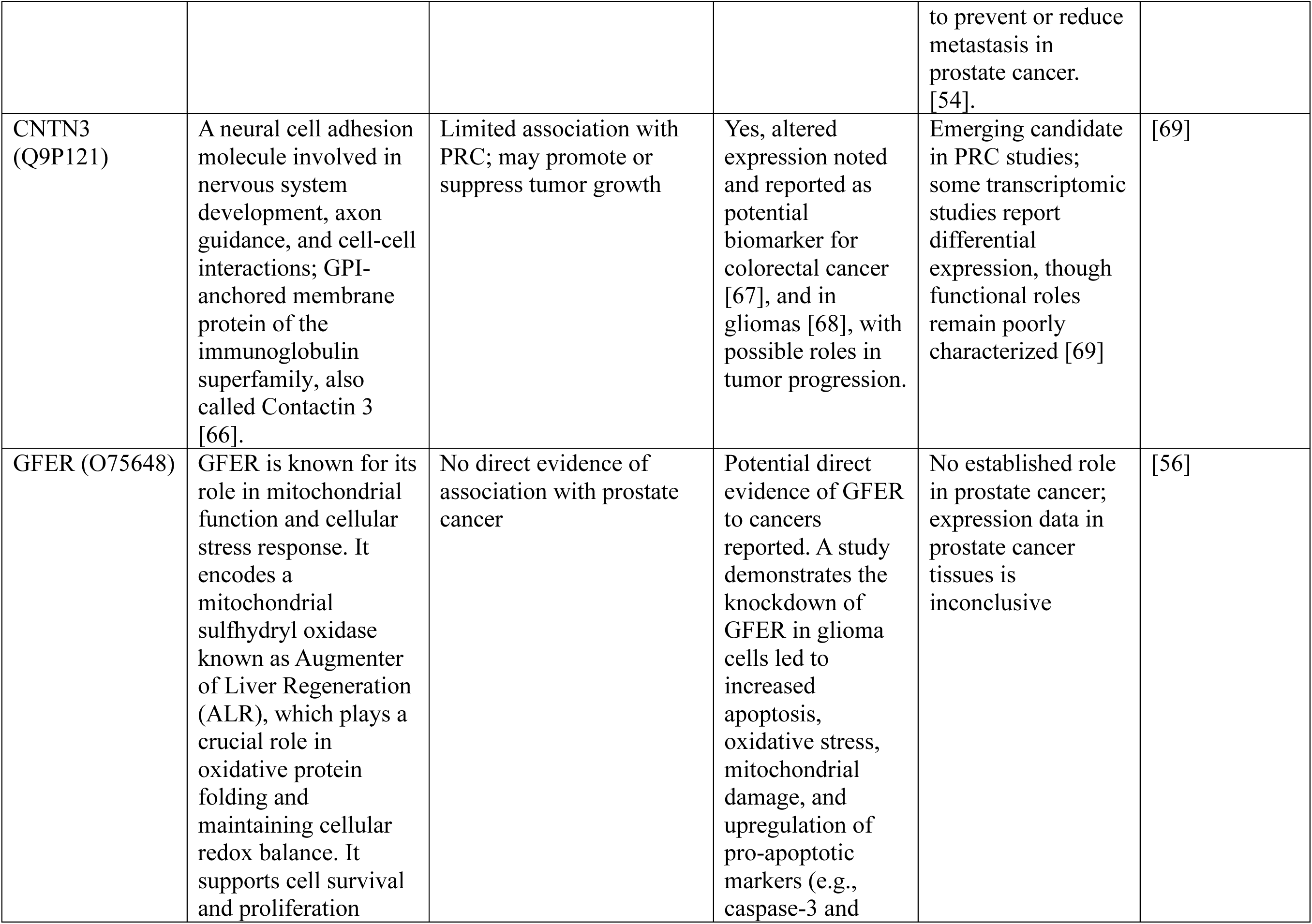
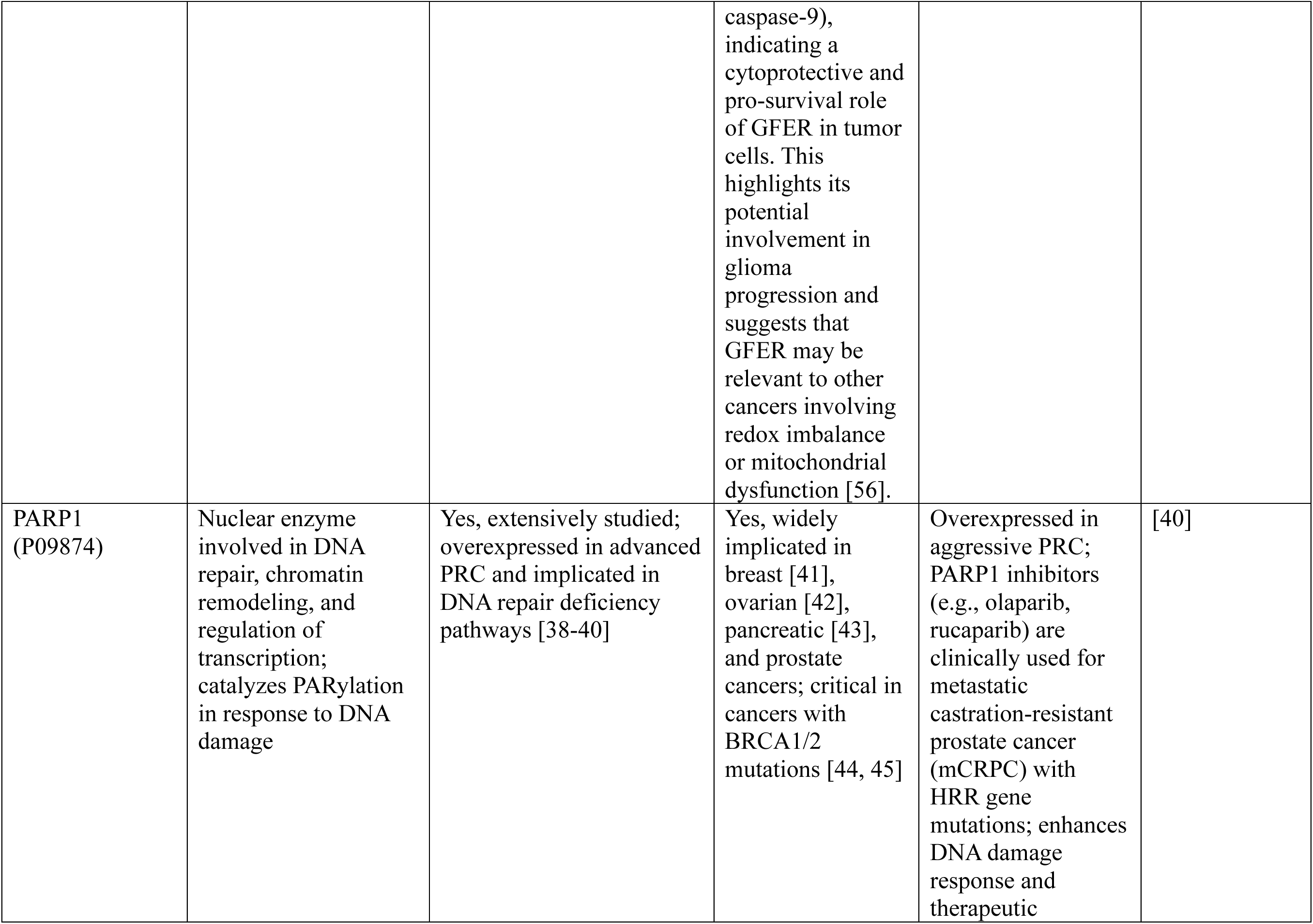
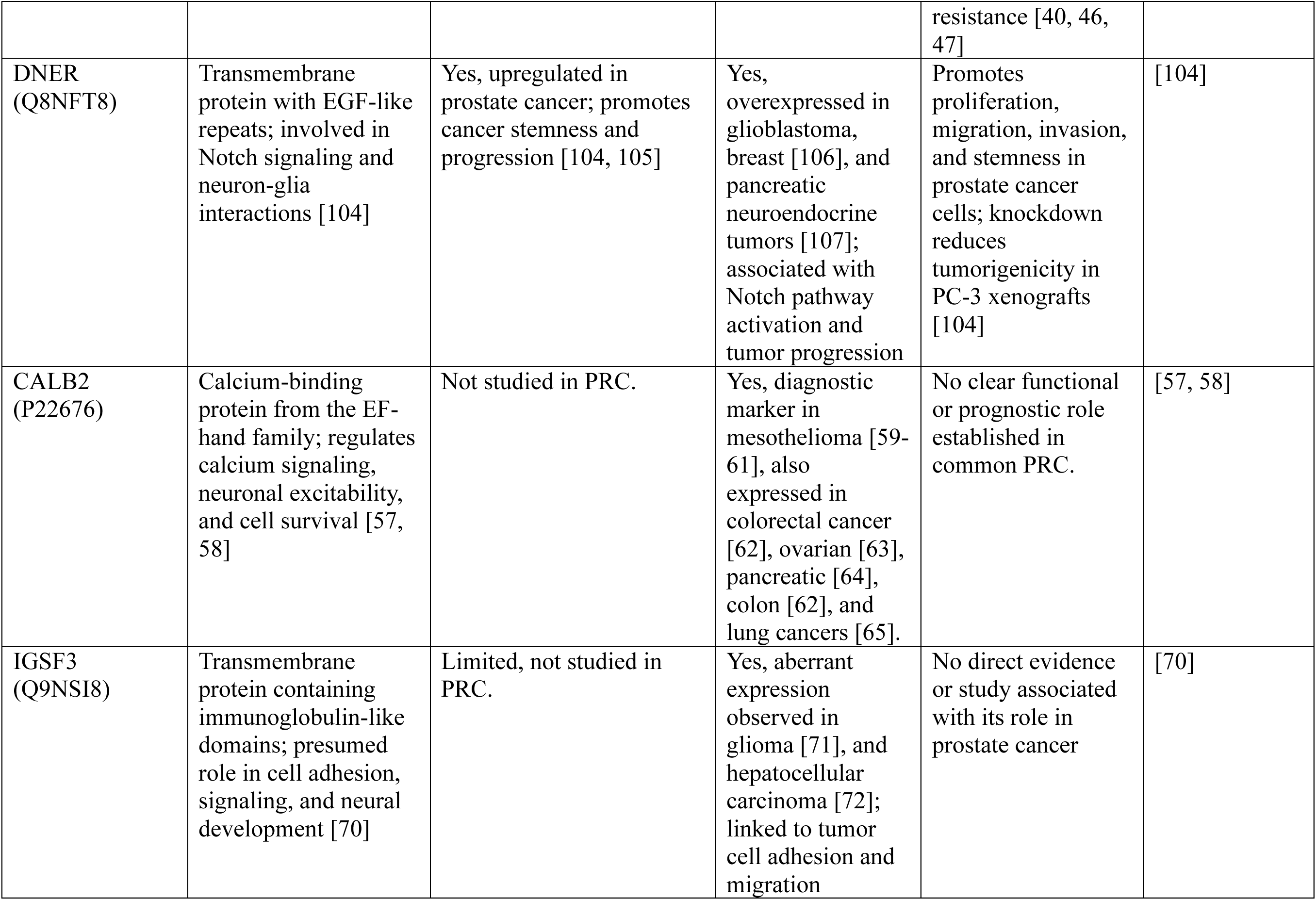
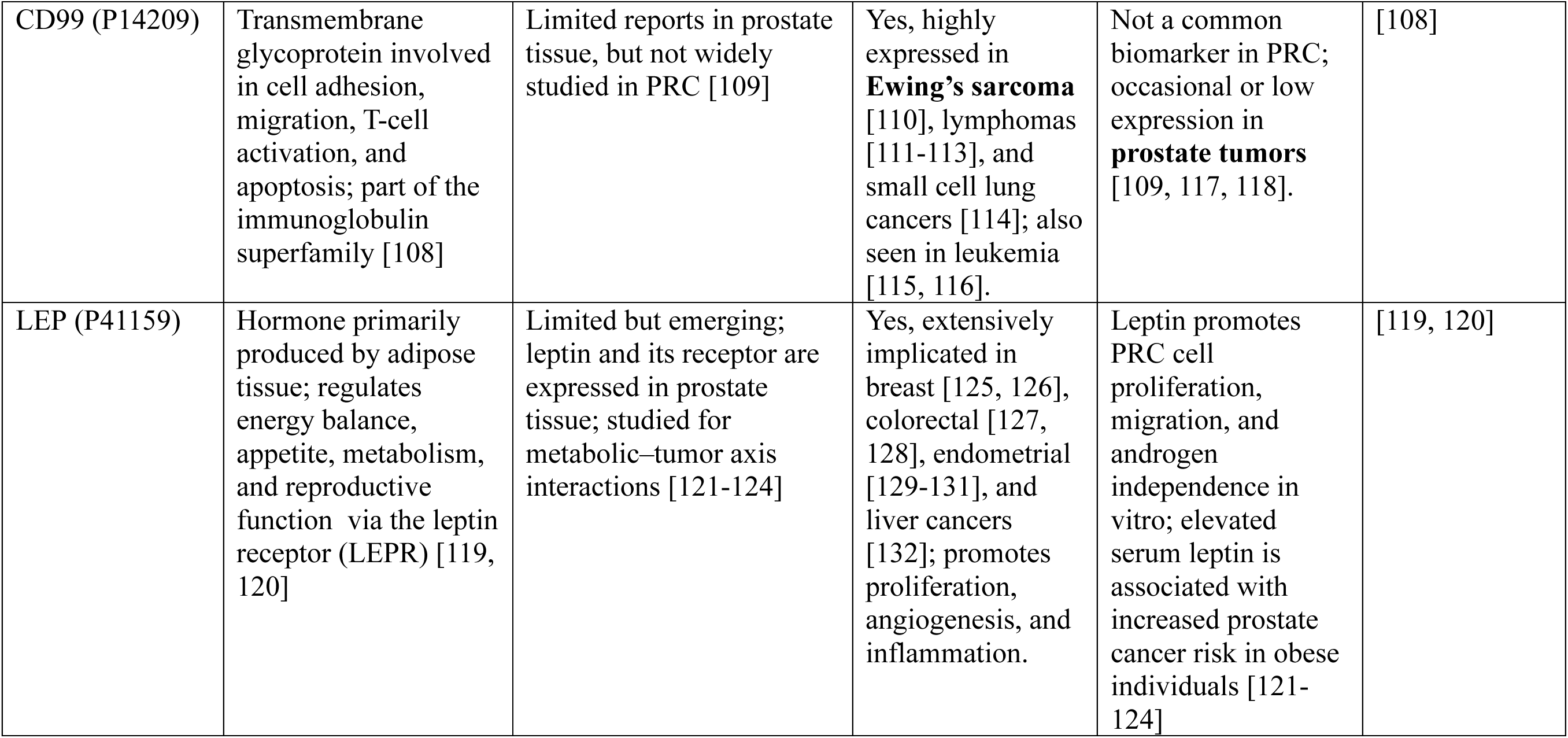
Functional Overview and Cancer Associations of GA Selected Proteins in Prostate Cancer (PRC). This table summarizes the biological functions, prior evidence in PRC, associations in other cancers, and hypothesized mechanistic roles in prostate tumorigenesis for each of the 14 proteins identified by the GA–LASSO pipeline.

Among the 14 proteins, MSMB, PARP1, FAP, COMP, DNER, and TNFRSF10A have strong literature support in the context of PRC. MSMB (beta-microseminoprotein) is consistently downregulated in PRC and has been widely proposed as a diagnostic and prognostic marker [26–37]. PARP1, involved in DNA damage repair, is overexpressed in aggressive PRC and targeted by approved therapies [38–47]. FAP and COMP, extracellular matrix components, are linked to tumor microenvironment remodeling and PRC progression [48–50], while TNFRSF10A is a pro-apoptotic receptor associated with androgen-independent PRC [51–53]. DNER is involved in apoptosis signaling and tumor cell adhesion and has been linked to mechanisms of resistance or metastasis in PRC [54, 55].

To assess the added value of the remaining 6 proteins not reported in literature to be involved in PRC, we trained classification models using only the eight literature-supported proteins. This restricted panel yielded lower classification performance (mean accuracy: 91.3%, ROC AUC: 0.97, F1: 0.91), compared to the full 14-protein panel (accuracy: 98.4%, ROC AUC: 0.99, F1: 0.98). These reductions were consistent across all metrics, including F1 scores, precision, and recall (Table 1). This highlights that the described approach identifies novel biomarkers which other methods of biomarker discovery cannot find. To contextualize these findings, we provide a structured overview of the molecular functions, prior cancer associations, and prostate cancer-specific roles of the 14 proteins (Table 2).

The decline in performance highlights the predictive contribution of the remaining six proteins: XG, GFER, PSPN, CNTN3, IGSF3, and CALB2. While these proteins are not widely characterized in PRC, several have been reported in other malignancies. For example, GFER is involved in mitochondrial redox homeostasis and has emerging roles in tumor growth and stress resistance [56]. PSPN, a neurotrophic factor, has been linked to aggressive neuroendocrine phenotypes [54]. CALB2 and CNTN3, though primarily studied in neurological contexts, have shown deregulation in cancers such as glioblastoma and colorectal cancer [57–69]. IGSF3, an immunoglobulin superfamily member, is associated with epithelial-mesenchymal transition and invasion in lung and pancreatic cancers [70–72]. XG, the most understudied of the group, is a GPI-anchored protein with potential implications in cell-cell interactions and immune recognition [73–76].

The inclusion of these proteins improved both sensitivity and specificity and enhanced class balance across all classification metrics. Importantly, their mechanistic interplay across apoptosis, extracellular matrix remodeling, calcium signaling, and neurotrophic pathways is illustrated in Figure 4, which integrates the GA-selected proteins into hallmark processes of prostate cancer progression. Their consistent recurrence across GA iterations, despite limited prior association with PRC, suggests that they may capture non-redundant, biologically meaningful variations related to tumor progression or microenvironmental dynamics. These results support the hypothesis that under-characterized proteins, when identified through data-driven approaches, can contribute significant diagnostic value and may highlight novel biological mechanisms. Overall, this analysis yields a compact and biologically coherent panel of plasma proteins capable of accurately distinguishing PRC from other tumor types, providing a strong basis for biomarker development and mechanistic follow-up studies.

**Figure 4:**
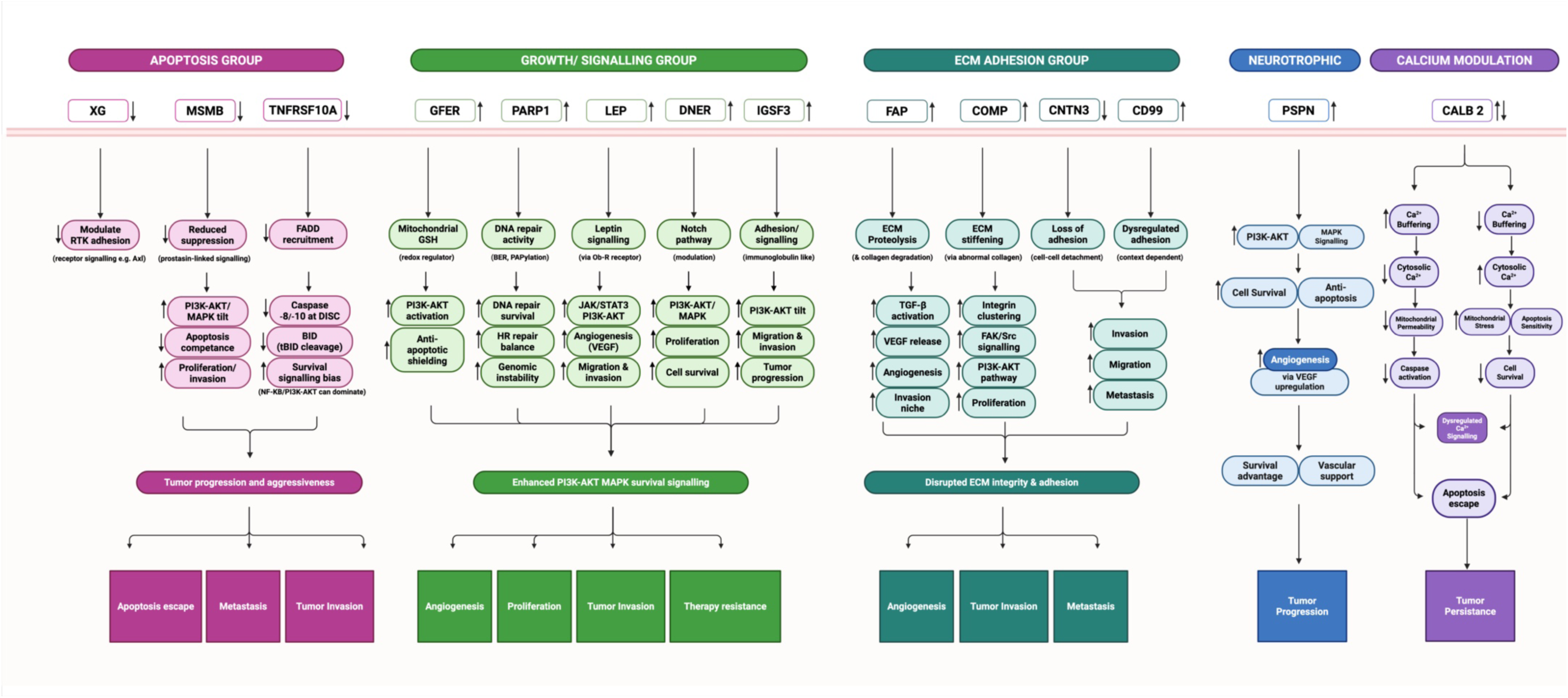
Mechanistic integration of GA-selected proteins into hallmark pathways of prostate cancer. The 14 proteins identified by the GA-LASSO pipeline cluster into five functional groups: apoptosis (XG, MSMB, TNFRSF10A), growth and survival signaling (GFER, PARP1, LEP, DNER, IGSF3), extracellular matrix adhesion and remodeling (FAP, COMP, CNTN3, CD99), neurotrophic signaling (PSPN), and calcium modulation (CALB2). Each group contributes to critical cancer hallmarks, including apoptosis escape, angiogenesis, proliferation, invasion, metastasis, and therapy resistance. The diagram illustrates how these proteins converge on PI3K-AKT/MAPK survival signaling and microenvironmental dysregulation, providing a mechanistic rationale for their combined predictive power in distinguishing prostate cancer.

## Discussion

We present a machine learning-based framework that identifies a minimal, high-performing plasma protein panel for prostate cancer (PRC) classification. The pipeline integrates a genetic algorithm (GA)-based protein selection with robust model training using bootstrap-aggregated LASSO logistic regression. Using a publicly available pan-cancer proteomic dataset, we demonstrate that compact and meaningful diagnostic signatures can be derived from small protein sets, and these panels yield superior classification performance compared to high-dimensional or purely statistical approaches.

The main contribution of this study is the GA-driven biomarker selection methodology. We performed 1,000 independent GA runs, each using a balanced dataset of PRC and non-PRC samples and a logistic regression fitness function optimized for ROC AUC. This iterative process consistently identified protein subsets of 25-30 proteins that yielded classification accuracies above 94%. By aggregating recurrence patterns across runs, we derived a ranked list and identified a 14-protein panel that achieved an average ROC AUC of 0.997, an F1 score of 0.980, and an accuracy of 98.0%, all with low variance across 100 bootstrapped replicates. Notably, performance plateaued beyond 14 proteins, confirming that the selected set was not only compact but also information-rich.

To benchmark our approach, we compared it directly to the published model by [12], in which glmnet-regularized classifiers were trained using 1,463 proteins. In that work, the full-protein models reached ∼90% accuracy, however, performance dropped substantially when the number of proteins was reduced. Notably, their top 14-protein panel achieved an ROC AUC of only 0.852. These comparisons emphasize that performance gains in our framework stem not only from biomarker selection but also from the robust classification strategy itself.

Across recent literature, prostate cancer classification models have shown AUCs typically ranging from 0.73 to 0.91, often using transcriptomic [133], cfDNA [134], flow cytometry [135], or multi-omics inputs. For example, studies using RNA-seq from TCGA cohorts or integrated transcriptomic panels have reported AUCs around 0.91-0.93, using up to 200 genes or combinations of clinical and omics data [133, 136]. Other approaches incorporating PSA kinetics, immune phenotyping, or DNA methylation achieved moderate sensitivity and specificity, often below 90% [137–139]. In contrast, our compact plasma protein panel exceeds these performances without the need for invasive sampling or complex multi-modal integration. This positions our framework as a promising foundation for blood-based, high-specificity diagnostic development.

Biological enrichment and literature annotation confirmed the plausibility of our findings. The 14-protein panel included both established and under-characterized proteins. While established biomarkers such as MSMB, PARP1, and FAP are well-recognized in PRC, less-characterized proteins including XG, IGSF3, and GFER, rarely discussed in the PRC literature, proved critical for achieving optimal classification. Importantly, models trained only on the eight well-established proteins from the panel showed a clear drop in performance, reinforcing the value of including novel, data-driven candidates.

To provide an integrative overview, we present a conceptual model that organizes the 14 proteins into functional axes driving prostate cancer progression (Figure 5). This conceptual framework underscores the biological coherence of the panel, linking statistical signal to mechanistic plausibility. Together, these results establish a machine learning framework capable of identifying small, interpretable, and biologically grounded diagnostic signatures. By integrating statistical analysis, evolutionary search, and literature-based validation, we provide a template for biomarker development that is both data-driven and biologically principled. The final 14-protein panel offers a promising basis for targeted assay development and provides a roadmap for extending this approach to other cancers and multi-omics platforms.

**Figure 5.**
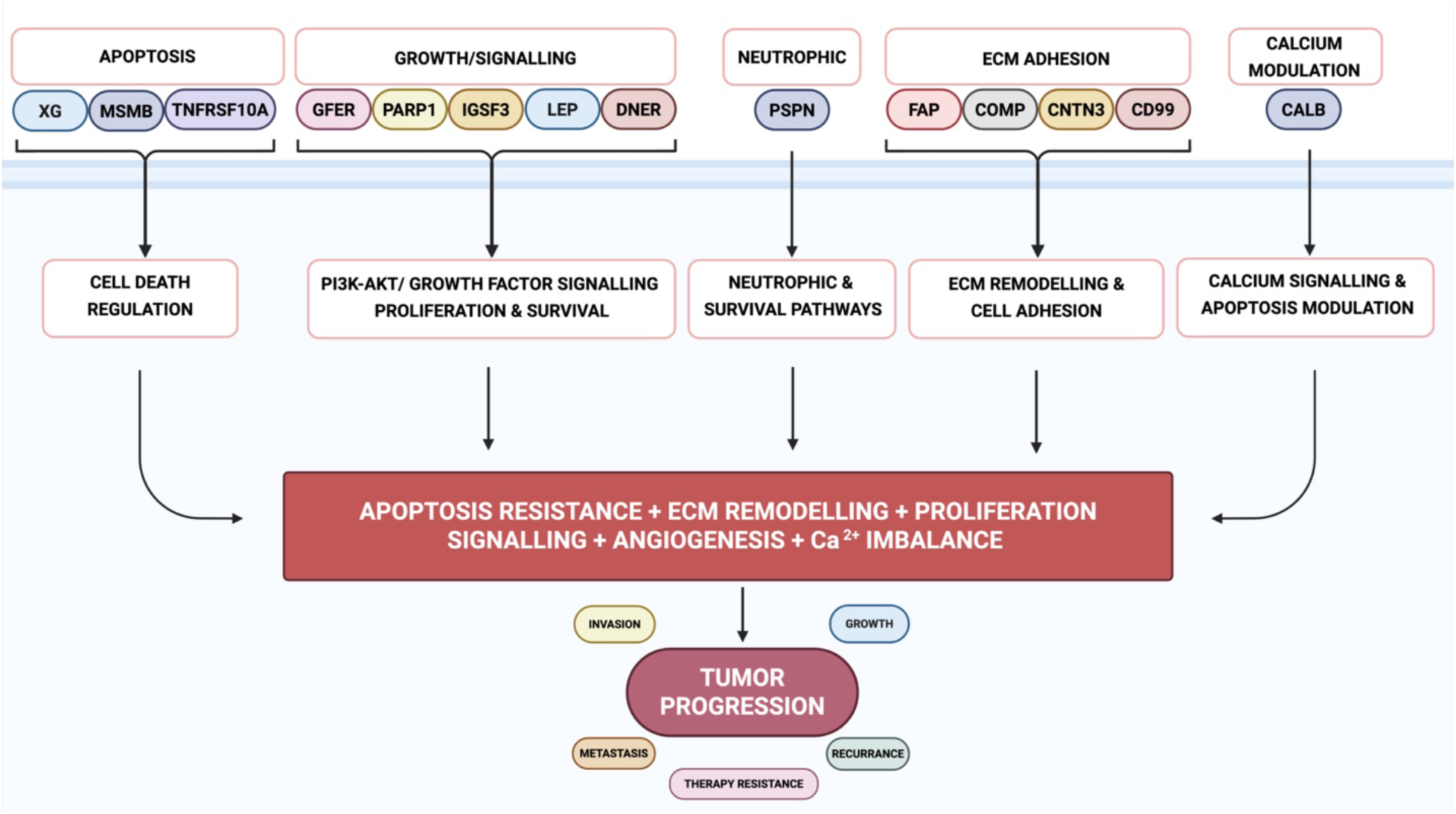
Conceptual model of GA-selected proteins in prostate cancer progression. The 14-protein panel is grouped into five functional categories: apoptosis (XG, MSMB, TNFRSF10A), growth/survival signaling (GFER, PARP1, LEP, DNER, IGSF3), extracellular matrix adhesion/remodeling (FAP, COMP, CNTN3, CD99), neurotrophic signaling (PSPN), and calcium modulation (CALB2). These pathways converge on apoptosis resistance, ECM remodeling, proliferative and angiogenic signaling, and Ca²⁺ imbalance, collectively driving tumor progression, invasion, metastasis, recurrence, and therapy resistance. This summary model highlights how under-characterized proteins complement well-established biomarkers in capturing key hallmarks of prostate cancer biology.

While the present study demonstrates the effectiveness of genetic algorithm-driven selection in identifying a compact, high-performing plasma protein panel for prostate cancer classification, certain limitations should be acknowledged. First, the analysis was based on a publicly available pan-cancer cohort rather than an independent, prospectively collected PRC-specific dataset. External validation using an independent cohort could not be performed due to the current unavailability of comparable Olink- or PEA-quantified prostate cancer proteomic datasets. Although extensive cross-validation and bootstrap aggregation were employed to minimize overfitting, future validation using independent plasma cohorts and clinical samples will be essential to confirm the generalizability of the findings. Additionally, functional characterization of underexplored proteins such as XG, GFER, and IGSF3 is warranted to elucidate their mechanistic roles in PRC biology. Future studies should aim to incorporate such experimental validation and assess clinical assay translation of the proposed 14-protein panel.

## Conclusion

This study presents a performance-optimized, machine learning-based pipeline that identifies a compact plasma protein signature for accurate prostate cancer prediction. Using genetic algorithm-driven protein selection and bootstrap-aggregated LASSO regression, we derived a 14-protein panel that consistently achieves over 98% accuracy and an ROC AUC of 0.99, outperforming previous models trained on much larger protein sets, offering a scalable, non-invasive alternative for clinical application. Unlike previous approaches based solely on differential expression or glmnet-based classification, our method directly optimizes for diagnostic performance and robustness. Importantly, we show that high accuracy can be achieved with minimal protein sets when guided by data-informed and performance-optimized selection strategies. While the final panel includes well-known markers like MSMB and PARP1, it also incorporates novel or under-characterized proteins that contribute unique diagnostic value, providing a base for its functional relevance in PRC biology. Functional enrichment and literature-based annotation confirm that the panel is not only statistically robust but also biologically meaningful. Compared to multi-omics or high-dimensional transcriptomic models, our compact, proteomics-only approach offers a scalable and clinically practical solution for early detection. This compact signature provides a strong foundation for future diagnostic assay development and highlights the value of combining biological insight with evolutionary optimization in biomarker discovery. Although this study was limited by the lack of independent validation owing to the absence of comparable Olink-based PRC proteomic datasets, the reproducible and data-driven framework established here provides a strong foundation for subsequent validation and clinical translation of the identified 14-protein panel.

## Statistical Analysis and Visualization

All statistical analyses were conducted in Python (v3.9) and R (v4.5.0). Machine learning metrics, including accuracy, precision, recall, F1 score, and ROC AUC, were computed using NumPy, pandas, and scikit-learn. LASSO logistic regression and genetic algorithm-based protein selection were implemented using scikit-learn and sklearn-genetic, respectively. Bootstrap resampling was used to assess performance stability across models. All visualizations were generated using matplotlib (v3.8) and seaborn (v0.13).

## Data Availability

Code and scripts for data analysis and visualization can be provided upon request.

## Author Contributions

S.A.S. and A.A.S. designed the study, S.A.S. preprocessed proteomic data, developed and implemented the GA-AutoML pipeline, conducted protein selection and classifier evaluation, performed biological interpretation, and wrote the manuscript. A.A.S. supervised the study, provided conceptual guidance on methodology and biological validation, and critically revised the manuscript. All authors reviewed and approved the final version of the manuscript.

## Acknowledgments

The authors gratefully acknowledge the University of Nizwa, Natural and Medical Sciences Research Center, for providing computational and infrastructural support.

